# A *mus-51* RIP allele for transformation of *Neurospora crassa*

**DOI:** 10.1101/043828

**Authors:** Zachary J. Smith, Stacy Bedore, Stephanie Spingler, Thomas M. Hammond

## Abstract

This report describes the construction and characterization of *mus-51^RIP70^*, an allele for high-efficiency targeted integration of transgenes into the genome of the model eukaryote *Neurospora crassa*. Two of the *mus-51^RIP70^* strains investigated in this work (RZS27.10 and RZS27.18) can be obtained from the Fungal Genetics Stock Center. The two deposited strains are, to our knowledge, genetically identical and neither one is preferred over the other for use in *Neurospora* research.

## Introduction

Non-homologous end joining, a DNA repair pathway that joins damaged DNA ends without regard for homology, was a hindrance to targeted transgene integration in *N. crassa* until Ninomiya *et al*. (2004) discovered that using an NHEJ mutant as a transformation host greatly increases the efficiency of this process. For example, one can achieve targeted transgene integration levels of nearly 100% by including either *mus-51*^*Δ*^*::hph* or *mus-51*^*Δ*^*::bar* in the transformation host's genetic background (Ninomiya *et al*., 2004; Colot *et al*., 2006). However, these alleles prevent one from using both *hph* and *bar* as selectable markers in consecutive transformations of the same host. Construction of a marker-free *mus-51* null allele could eliminate this deficiency. Below, we describe the construction and characterization of a *mus-51* null allele called *mus-51*^*RIP70*^. Additionally, we describe a method to distinguish between *mus-51*^+^ and *mus-51*^*RIP70*^ genotypes by restriction endonuclease-mediated digestion of PCR products.

## Materials and Methods

### Strains and growth conditions

Vogel’s minimal medium (VMM) (Vogel, 1956) with or without histidine (0.5 g/L) was used for vegetative cultures. Synthetic Crossing Medium (SCM) (Westergaard and Mitchell, 1947) with or without histidine (0.5 g/L) was used for crosses. BDS medium (1% sorbose, 0.05% glucose, and 0.05% fructose) (Brockman and de Serres, 1963) with or without methyl methanesulfonate (MMS, 0.22 μl/ml) was used in a preliminary screen for *mus-51*^*RIP*^ alleles. BDS medium with hygromycin (200 μg/ml) and/or cyclosporin A (5 μg/ml) was used to screen for hygromycin and cyclosporin A resistance. Top and Bottom Agar were used in transformation experiments as previously described (Harvey *et al*., 2014). The key strains used in this study are listed in Table 1.

**Table 1.** Strains used in this study.

| Strain Name | Genotype |
| --- | --- |
| FGSC 6103 | *his-3 A* |
| FGSC 9716 | *his-3 a* |
| HSS1.21.4 | *his-3<sup>+</sup>::mus-51<sup>2170</sup> A* |
| RSB1.70 | *his-3<sup>+</sup>::mus-51<sup>2170</sup>; mus-51<sup>RIP70</sup> A* |
| F2-26 | *rid; fl a* |
| RZS27.4 | *rid a* |
| RZS27.6 | *rid a* |
| RZS27.10 | *rid; mus-51<sup>RIP70</sup> a* |
| RZS27.18 | *rid; mus-51<sup>RIP70</sup> a* |
| P8-65 | *csr-1<sup>Δ</sup>::hph a* |
| P6-07 | *rid A* |

### Plasmid construction

Plasmid pTH1256.1 was constructed by cloning the *hph* selectable marker from pCB1004 (Carroll *et al*., 1994) into the *ApaI* site of pBM61 (Margolin *et al*., 1997). Oligonucleotides P518 and P519 (Table 2) were then used to PCR-amplify a 2170 base pair (bp) fragment of *mus-51*^+^. This PCR product was cloned into the *Not*I site of plasmid pTH1256.1 to create plasmid pSS2.12. Plasmid pSS2.12 thus contains the sequences necessary to insert the 2170 bp *mus-51*^+^ fragment next to the *his-3* locus while converting *his-3* to *his-3*^+^.

**Table 2.**
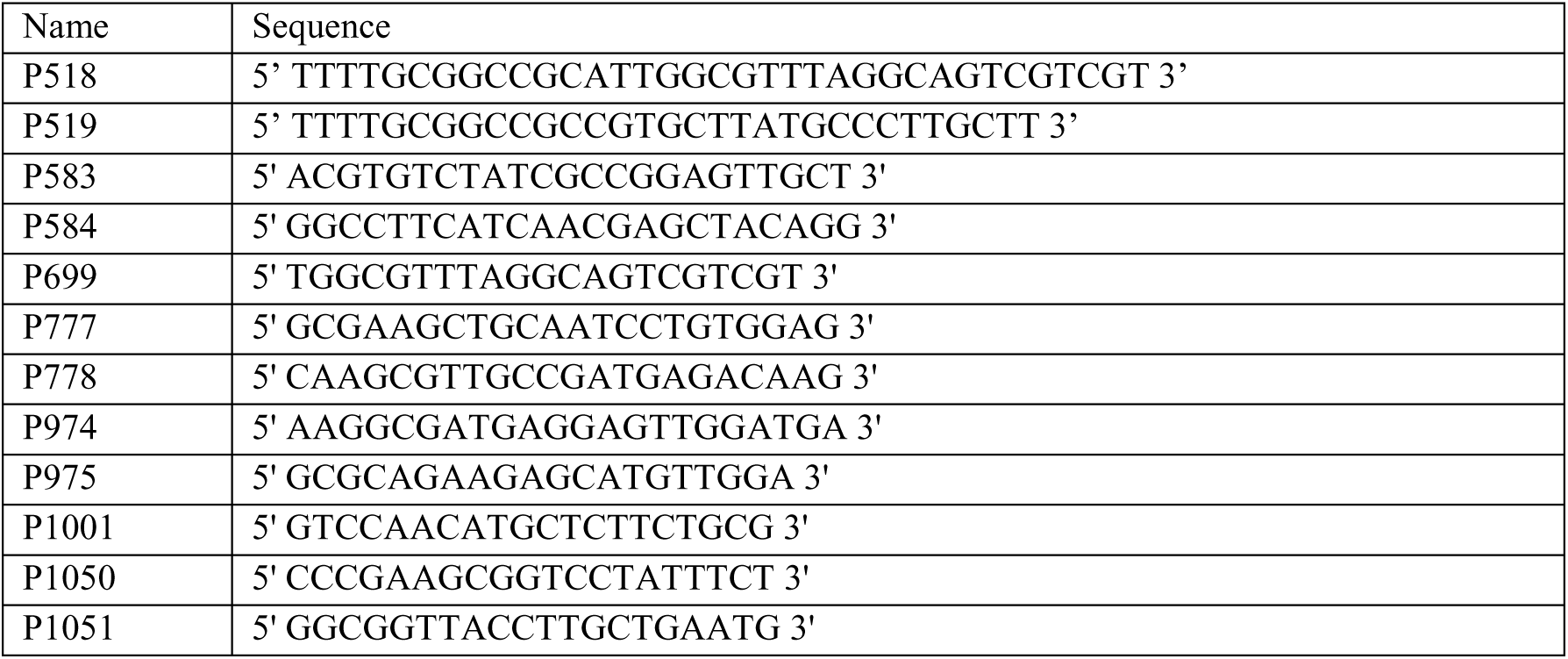
Oligonucleotides used in this study.

### Transformation of *N. crassa*

Plasmid pSS2.12 was linearized with the restriction endonuclease *SspI* and transformed into FGSC 6103 by electroporation with selection for histidine prototrophy as previously described (Margolin *et al*., 1997). The homokaryotic strain HSS1.21.4 was isolated from a heterokaryotic transformant with a microconidium-based homokaryon isolation technique (Ebbole and Sachs, 1990).

### Isolation of *mus-51*^*RIP70*^

HSS1.21.4 was crossed with FGSC 9716 by simultaneous inoculation of each strain to opposite sides of a 100 mm petri dish containing SCM plus histidine. The petri dish was incubated on a laboratory bench top for five weeks. Ascospores were collected, soaked in sterile water at 4 °C for over 24 hours, heat-shocked at 60 °C for 30 minutes, and plated onto VMM. Histidine-prototrophs were selected and screened for sensitivity to methyl methanesulfonate (MMS). The *mus-51* coding region of an MMS-sensitive progeny named RSB1.70 was analyzed by Sanger sequencing, found to be mutated, and named *mus-51*^*RIP70*^. Next, protoperithecia of F2-26 were fertilized with conidia from RSB1.70. This cross produced progeny RZS27.4, RZS27.6, RZS27.10, and RZS27.18. The *mus-51* locus in each of these four strains was PCR-amplified with oligonucleotides P777 and P778 and the PCR products were sequenced by Sanger sequencing with oligonucleotides P699, P974, P975, P1001, P1050, and P1051. Sequences were analyzed with Bioedit 7.2.5 (Hall, 1999). The full sequence of *mus-51*^*RIP70*^ can be obtained from GenBank under accession number KU860571.

### Polymerase chain reaction (PCR)

Genomic DNA was isolated from lyophilized mycelia with the IBI Scientific Plant Genomic DNA Mini Kit. PCR was performed with New England Biolabs Phusion High-Fidelity DNA polymerase or MidSci Bullseye Taq DNA polymerase. When restriction endonuclease-mediated digestion of PCR products was needed to differentiate between two products of similar size, 2.5 μl of a completed PCR reaction was digested with a restriction endonuclease in a 25 μl reaction under standard conditions.

### The *csr-1*^+^ gene deletion assay

A 3718 bp *csr-1*^*+*^ deletion vector was obtained by PCR-amplifying the *csr-1*^*Δ*^*::hph* locus from strain P8-65 with oligonucleotides P583 and P584. The PCR product was purified with an IBI scientific Gel/PCR DNA Fragment Extraction Kit and 500 ng were electroporated into conidia as previously described (Margolin *et al*., 1997) with selection for hygromycin resistance. Hygromycin-resistant transformants were then screened for resistance to cyclosporin A.

## Results and Discussion

### The *mus-51*^*RIP70*^ allele is 84% identical to wild type

Repeat-induced point mutation (RIP) introduces transition mutations into repeated DNA sequences within the nuclei of sexual cells just prior to meiosis (Cambareri *et al*., 1989; Selker, 1990). Therefore, we used RIP in an attempt to generate a *mus-51* null allele at its native location on chromosome IV by first placing a 2170 bp fragment of *mus-51+* next to the *his-3*^*+*^ locus on chromosome I. We then placed the resulting *his-3*^+^*::mus-51^2170^*-carrying transformant (HSS1.21.4) through a sexual cross with strain FGSC 9716. Strains mutated in *mus-51* are sensitive to the DNA damaging agent methyl methanesulfonate (MMS) (Ninomiya *et al*., 2004). We thus selected RSB1.70, an MMS sensitive-progeny of HSS1.21.4 × FGSC 9716 (data not shown), for further analysis. Sanger sequencing confirmed RSB1.70 to carry a mutated *mus-51* allele, which we named *mus-51*^*RIP70*^.

The *mus-51*^*RIP70*^ allele contains 341 transition mutations spread over a 2134 bp region of chromosome IV (Figure 1). The mutations begin 211 bp before the start of the *mus-51*^*+*^ coding sequence and they end 255 bp before the end of the coding sequence (Figure 1). If we assume that all RIP mutations result from C to T transition events, 228 mutations must have originated on the coding strand and 113 mutations must have originated on the template strand (Figure 1, red lines). The high number of mutations suggests that *mus-51*^*RIP70*^ is a null allele. Moreover, the *mus-51*^*RIP70*^ allele encodes 30 early stop codons and over 100 amino acid substitutions relative to *mus-51*^+^.

**Figure 1:**
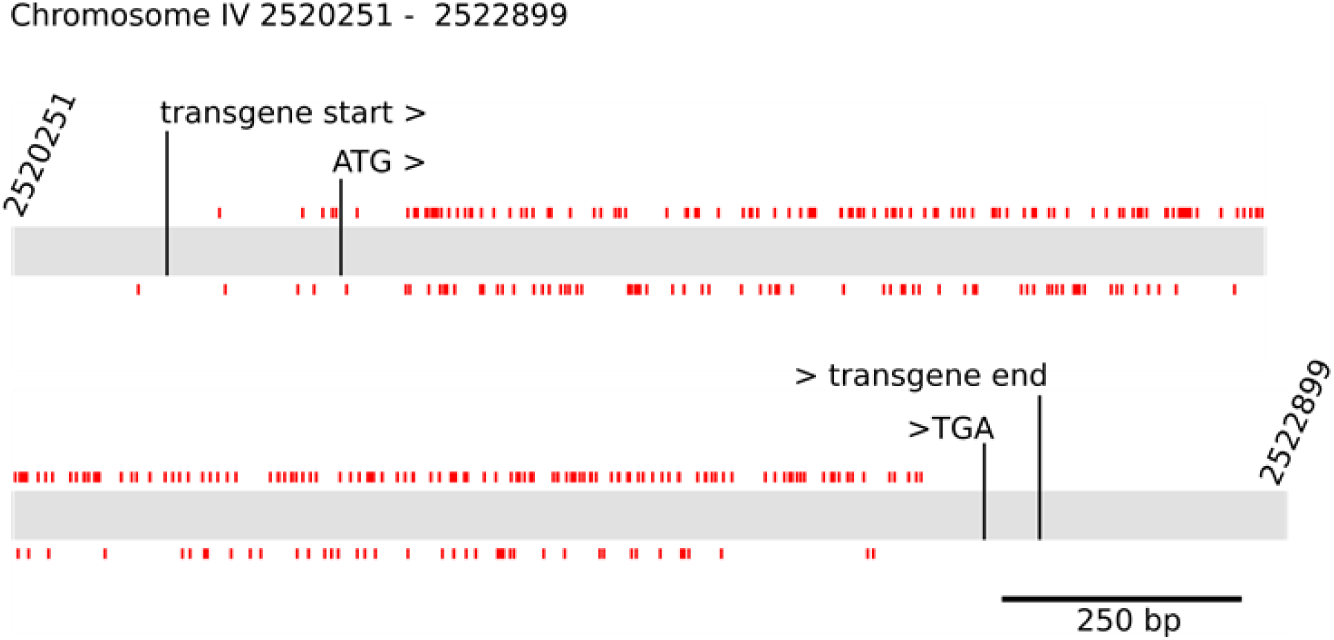
The *mus-51*^*RIP70*^ allele contains 341 transition mutations. Positions 2,520,251 through 2,522,899 on chromosome IV of the *N. crassa* reference genome (version 12) are depicted in the diagram. The 2170 bp fragment, which was inserted next to the *his-3*^*+*^ locus on chromosome I, is delineated by black vertical lines labeled "transgene start >" and "> transgene end". The coding sequence of *mus-51*^*+*^ is marked with the black vertical lines labeled "ATG>" and ">TGA". The short vertical red lines denote the locations of C to T transitions on the coding strand (shown above chromosome) or the template strand (shown below chromosome).

### The *mus-51*^*RIP70*^ allele increases the efficiency of targeted transgene integration

The efficiency of targeted transgene integration when using *mus-51*^*RIP70*^ in the genetic background of a transformation host was measured with a csr-1+-gene deletion assay. Deletion of *csr-1*^*+*^ enhances resistance to cyclosporin A (Bardiya and Shiu, 2007), allowing one to identify *csr-1*^*Δ*^ genotypes by screening on medium containing the compound. We transformed four strains, RZS27.4 (*mus-51^+^*), RZS27.6 (*mus-51^+^*), RZS27.10 (*mus-51*^*RIP70*^), and RZS27.18 (*mus-51*^*RIP70*^), with a *csr-1*^*Δ*^*::hph* deletion vector. While nearly all *mus-51*^*RIP70*^ transformants (99.5%) were resistant to cyclosporin A, only 37.5% of the *mus-51^+^* transformants were resistant (Figure 2). To confirm that cyclosporin A resistance resulted from replacement of *csr-1*^+^ allele by the *csr-1*^*Δ*^*::hph* deletion vector, PCR was used to examine the *csr-1* locus in a cyclosporin A-resistant and a cyclosporin A-susceptible transformant from each transformation host. While all of the cyclosporin A-resistant isolates carried the *csr-1*^*Δ*^*::hph* allele at the *csr-1* locus, none of the susceptible isolates did (Figure 3). These results confirm that the *mus-51*^*RIP70*^ allele can be used for high-efficiency targeted integration of transgenes in *N. crassa*.

**Figure 2:**
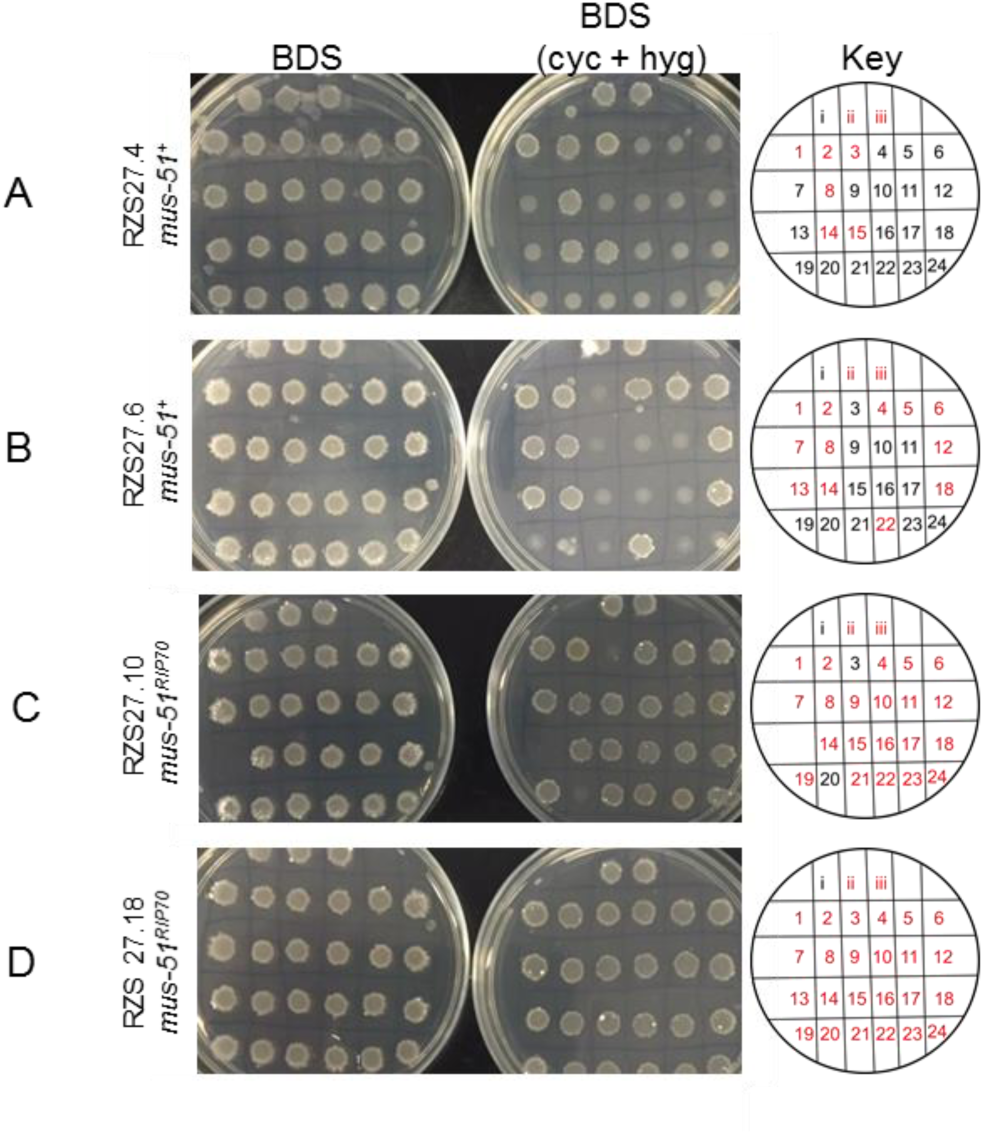
The *mus-51*^*RIP70*^ allele improves the efficiency of transgene integration by homologous recombination. A *csr-1*^*^+^*^ deletion assay was performed with two *mus-51*^+^ strains (RZS27.4 and RZS27.6) and two *mus-51*^*RIP70*^ strains (RZS27.10 and RZS27.18). Each strain was transformed with a *csr-1*^+^ deletion vector. The *csr-1*^*Δ*^*::hph* deletion vector replaces the entire coding region, as well as upstream regulatory sequences, with an *hph* selectable marker (Bardiya and Shiu, 2007). Only transformants resulting from replacement of *csr-1*^+^ with *csr-1*^*Δ*^*::hph* should be resistant to cyclosporin A. A total of 96 transformants were isolated, 24 for each of the transformation hosts. Conidial suspensions were prepared from each transformant and tested for growth on minimal BDS medium with or without hygromycin and cyclosporin (please note that hygromycin was an unnecessary addition because all transformants were previously shown to be resistant to hygromycin). Cyclosporin A resistance is indicated by the formation of large dense colonies with rough borders, while susceptibility is indicated by small thin colonies with smooth borders. A) A total of 6 out of 24 RZS27.4 transformants were resistant. B) A total of 12 out of 24 RZS27.6 transformants were resistant. C) A total of 21 out of 23 RZS27.10 transformants were resistant. Strain 13 did not grow on minimal medium so it was not included. D) A total of 24 out of 24 RZS27.18 transformants were resistant. Key: i) P6-07, ii) P8-65, iii) P6-07 and P8-65 [*i.e*., a suspension of conidia from both strains]. Numbers 1-24 refer to the 24 transformants from each transformation host. Red font is used to indicate resistance to cyclosporin A.

**Figure 3:**
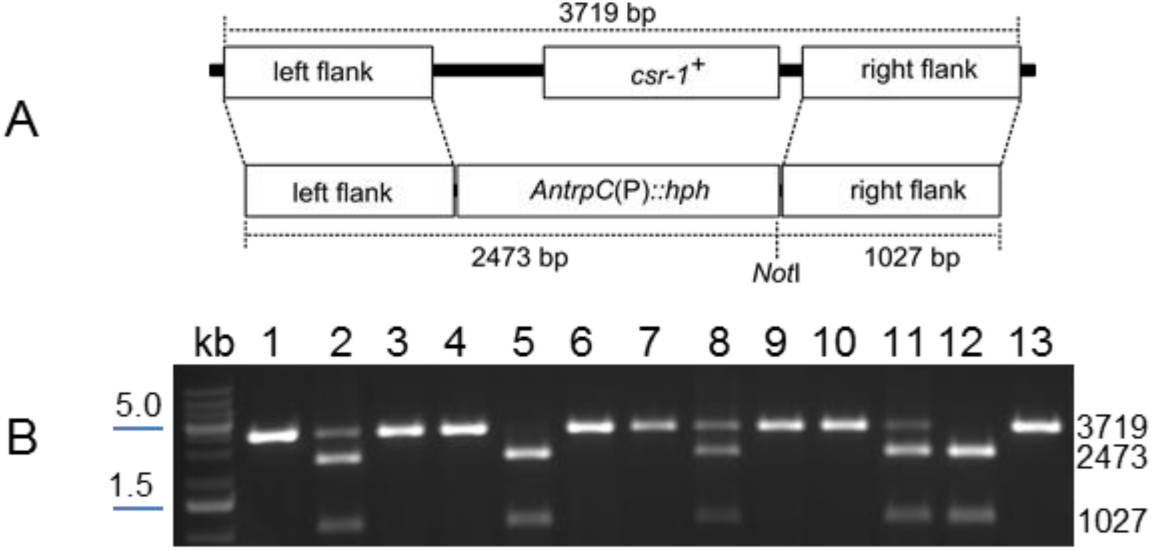
PCR analysis confirms that resistance to cyclosporin A is due to replacement of the *csr-1*^+^ allele with *csr-1*^*Δ*^*::hph*. A) A diagram of the *csr-1*^+^ locus (top) relative to the *csr-1*^*Δ*^*::hph* vector (bottom). Oligonucleotides P583 and P584 amplify a 3719 bp product from the *csr-1* locus when it contains the *csr-1*^+^ allele and a 3500 bp PCR product from the locus when it contains the *csr-1*^*Δ*^*::hph* allele. The restriction endonuclease NotI can be used to distinguish between these two PCR products of similar length. While NotI will digest a *csr-1*^*Δ*^*::hph* PCR product into two fragments of 2473 and 1027 bp, it should not digest a PCR product from *csr-1*^+^. B) Eleven transformants from the *csr-1*^^+^^ gene deletion assay were analyzed by PCR. The four host strains (host), a cyclosporin A-resistant transformant (CART) from each host, and a cyclosporin A-susceptible transformant (CAST) from each host (if available) were analyzed. The *csr-1* locus in each strain was PCR amplified with oligonucleotides P583 and P584, digested with NotI, and analyzed by gel electrophoresis. Only CART strains (lanes 2, 5, 8, 11) and the positive control P8-65 (lane 12) produced PCR products with the expected pattern for the *csr-1*^*Δ*^*::hph* allele. The transformants were not purified by homokaryon isolation, thus *csr-1*^+^ and *csr-1*^*Δ*^*::hph* specific patterns in some lanes is expected (*e.g*., lanes 2, 8, and 11). The 5.0 and 1.5 kb positions, according to the included DNA ladder, are indicated to the left of the gel image. Lane information: kb) DNA ladder; 1) RZS27.4, host; 2) RZS27.4, CART; 3) RZS27.4, CAST; 4) RZS27.6, host; 5) RZS27.6, CART; 6) RZS27.6, CAST; 7) RZS27.10, host; 8) RZS27.10, CART; 9) RZS27.10, CAST; 10) RZS27.18, host; 11) RZS27.18, CART; 12) P8-65, *csr-1*^*Δ*^*::hph* control; and 13) F2-26, *csr-1*^+^ control.

### The restriction endonuclease *Rsa*I can distinguish between *mus-51*^*RIP70*^ and *mus-51*^+^ alleles

One disadvantage of the *mus-51*^*RIP70*^ allele is the inability to identify the allele by screening for growth on a common antibiotic. To address this issue, we devised a simple PCR-assay to distinguish between *mus-51*^*RIP70*^ and *mus-51*^+^ alleles. In this assay, the *mus-51* locus is PCR-amplified with oligonucleotides P974 and P975 and the resulting 478 bp PCR product is digested with the restriction endonuclease *Rsa*I. The digested product is then analyzed by standard agarose gel electrophoresis. RsaI will only digest the *mus-51*^+^ PCR product (Figure 4). This procedure can also be combined with the conidial-PCR method of Henderson et al., (2005) for increased efficiency.

**Figure 4:**
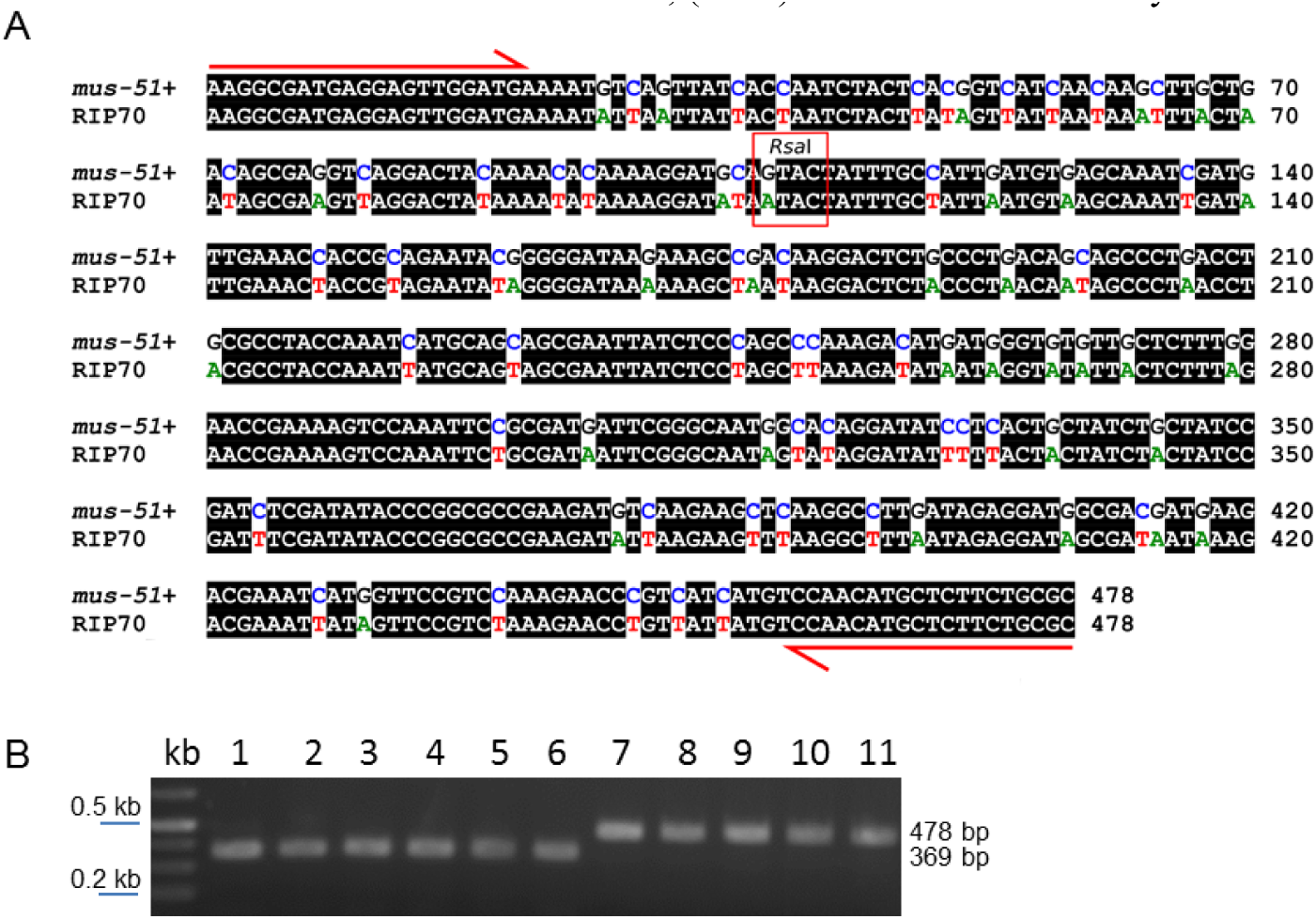
The *mus-51*^*RIP70*^ allele can be distinguished from *mus-51*^+^ by RsaI-based digestion of PCR products. A) Partial DNA sequences for *mus-51*^+^ and *mus-51*^*RIP70*^ are shown. Note that oligonucleotides P974 and P975 anneal perfectly to both alleles (red arrows). Although these primers amplify a PCR product of identical length from *mus-51*^+^ and *mus-51*^*RIP70*^, an RsaI recognition site (5' GTAC 3'), which exists in *mus-51*^+^ but not *mus-51*^*RIP70*^ (red rectangle), can be used to distinguish the PCR products. B) Oligonucleotides P974 and P975 were used to PCR-amplify the *mus-51* region from the 11 strains described in Figure 3B. The PCR products were digested with RsaI and analyzed by agarose gel electrophoresis. The *mus-51*^+^ allele produces two fragments with sizes of 369 and 109 bp, but the 109 bp fragment is often too faint to detect without additional gel staining. The *mus-51*^*RIP70*^ allele produces a single fragment of 478 bp. Lanes are labeled as in Figure 3B.

## Acknowledgements

We thank the fungal genetics stock center for strains FGSC 6103 and FGSC 9716 and Dr. Patrick Shiu for strains P6-07, P8-65, and F2-26. We are grateful to members of the Hammond Lab for technical assistance. This work was supported by a grant from the National Institute of Child Health and Human Development (NICHD, 1R15HD076309-01).

## References

Bardiya, N., and Shiu, P. K. T. 2007. Cyclosporin A-resistance based gene placement system for Neurospora crassa. Fungal Genet. Biol. 44:307–314.

Brockman, H. E., and de Serres, F. J. 1963. Sorbose Toxicity in Neurospora. Am. J. Bot. 50:709–714.

Cambareri, E. B., Jensen, B. C., Schabtach, E., and Selker, E. U. 1989. Repeat-induced G-C to A-T mutations in Neurospora. Science 244:1571–1575.

Carroll, A. M., Sweigard, J. A., and Valent, B. 1994. Improved vectors for selecting resistance to hygromycin. Fungal Genet. Newsl. 41:22.

Colot, H. V., Park, G., Turner, G. E., Ringelberg, C., Crew, C. M., Litvinkova, L., Weiss, R. L., Borkovich, K. A., and Dunlap, J. C. 2006. A high-throughput gene knockout procedure for Neurospora reveals functions for multiple transcription factors. Proc. Natl. Acad. Sci. U. S. A. 103:10352–10357.

Ebbole, D., and Sachs, M. 1990. A rapid and simple method for isolation of Neurospora crassa homokaryons using microconidia. Fungal Genet. Newsl. 37:17–18.

Hall, T. A. 1999. BioEdit: a user-friendly biological sequence alignment editor and analysis program for Windows 95/98/NT. Nucleic Acid Symp Ser 41:95–98.

Harvey, A. M., Rehard, D. G., Groskreutz, K. M., Kuntz, D. R., Sharp, K. J., Shiu, P. K. T., and Hammond, T. M. 2014. A critical component of meiotic drive in Neurospora is located near a chromosome rearrangement. Genetics 197:1165-1174.

Margolin, B. S., Freitag, M., and Selker, E. U. 1997. Improved plasmids for gene targeting at the his-3 locus of Neurospora crassa by electroporation. Fungal Genet. Newsl. 44:34–36.

McCluskey, K., Wiest, A., and Plamann, M. 2010. The Fungal Genetics Stock Center: a repository for 50 years of fungal genetics research. J. Biosci. 35:119–126.

Ninomiya, Y., Suzuki, K., Ishii, C., and Inoue, H. 2004. Highly efficient gene replacements in Neurospora strains deficient for nonhomologous end-joining. Proc. Natl. Acad. Sci. U. S. A. 101:12248–12253.

Selker, E. U. 1990. Premeiotic instability of repeated sequences in Neurospora crassa. Annu. Rev. Genet. 24:579–613.

Henderson, S. T., Eariss, G. A., and Catcheside, D. E. A. 2005. Reliable PCR amplification from Neurospora crassa genomic DNA obtained from conidia. Fungal Genet. Newsl. 52:24.

Vogel, H. J. 1956. A convenient growth medium for Neurospora (Medium N). Microb. Genet. Bull. 13:42–43.

Westergaard, M., and Mitchell, H. K. 1947. Neurospora V. A synthetic medium favoring sexual reproduction. Am. J. Bot. 34:573–577.

